# SEPIA: Simulation-based Evaluation of Prioritization Algorithms

**DOI:** 10.1101/2020.11.23.394890

**Authors:** Kimberly Almaraz, Tyler Jang, McKenna Lewis, Titan Ngo, Miranda Song, Niema Moshiri

## Abstract

**Background:** The ability to prioritize people living with HIV by risk of future transmissions could aid public health officials in optimizing epidemiological intervention. While methods exist to perform such prioritization based on molecular data, their effectiveness and accuracy are poorly understood, and it is unclear how one can directly compare the accuracy of different methods. We introduce SEPIA (Simulation-based Evaluation of PrIoritization Algorithms), a novel simulation-based framework for determining the effectiveness of prioritization algorithms. Under several metrics of effectiveness that we propose, we utilize various properties of the simulated contact networks and transmission histories to compare existing prioritization approaches: one phylogenetic (ProACT) and one distance-based (growth of HIV-TRACE transmission clusters).

**Results:** Using all metrics of effectiveness that we propose, ProACT consistently slightly outperformed the transmission cluster growth approach. However, both methods consistently performed just marginally better than random, suggesting that there is significant room for improvement in prioritization tools.

**Conclusion:** We hope that, by providing ways to quantify the effectiveness of prioritization methods in simulation, SEPIA will aid researchers in developing novel tools for prioritizing people living with HIV by risk of future transmissions.

## Background

Molecular data gathered on human immunodeficiency virus (HIV) is useful for understanding the systems of epidemic spread of HIV. Such understanding can better allow us to intervene and treat high-risk groups of individuals. Methods of epidemic intervention include treatments such as antiretroviral therapy (ART) and awareness programs [1]. When people living with HIV adhere to ART, they may experience viral suppression, which results in a significantly reduced risk of transmission. Thus, ART distribution and adherence is a potentially effective approach to combating the spread of HIV. However, a major issue for public health officials is the limited amount of resources available. Because of this, officials need to ensure that their limited resources are allocated optimally to best reduce the spread of an epidemic.

In many parts of the world, when testing and treating individuals living with HIV, it has become standard practice to record various pieces of metadata about the patient, including a viral genomic sequence (often of the *pol* and *gag* regions). This information is often used to determine groups of individuals with a high risk of future transmission, which can allow public health officials to better allocate their limited resources [2]. The prioritization of people living with HIV can be explored in a computational framework: given a list of individuals along with metadata and a viral sequence from each individual, order the individuals in descending order of inferred risk of future transmission.

Molecular epidemiology provides a natural framework for prioritizing individuals from viral sequence data. Currently, the standard approach is to use HIV-TRACE [3] to infer transmission clusters based on pairwise distances between sequences, monitor the growth of the transmission clusters over time, and prioritize individuals in descending order of transmission cluster growth. ProACT [4], on the other hand, is a prioritization approach that utilizes properties of a phylogeny inferred from the viral sequences.

The following questions naturally arise: how well does a given prioritization method perform, and which method is superior in specific contexts? With real-world data, the ground truth of who transmitted to whom is typically unavailable or error-prone. Further, even *with* a known transmission history, it is unclear how to quantify effectiveness: do we count the number of transmissions from a single individual? Do we count the total number of transmissions in a transmission chain seeded from a single individual? Rather than the transmission network itself, perhaps we are interested in properties of the underlying contact network (e.g. prioritize individuals with large numbers of social contacts)? Thus, it is unclear how to even quantitatively assess the performance of different prioritization methods.

To address this open problem, we introduce SEPIA (Simulation-based Evaluation of PrIoritization Algorithms), a novel simulation-based framework for measuring the effectiveness of prioritization algorithms. SEPIA utilizes simulated epidemic data, such as those generated by FAVITES [5] or PANGEA.HIV.sim [6], to define a ground truth with which prioritization methods can be directly compared. The user runs their prioritization method on a simulated dataset, and then given the prioritization as well as the simulated dataset, SEPIA will measure the effectiveness of the prioritization using any of the metrics defined below.

## Methods

Given a prioritization, SEPIA computes an effectiveness score according to one of several metrics we have implemented, defined below.

- **Metric 1: Direct Transmissions** This metric aims to quantify the direct impact of each individual *u* on the spread of the virus within a population by counting the total number of individuals to whom *u* directly transmitted.
- **Metric 2: Transmission Rate** This metric aims to quantify the rate of transmission of each individual *u*, giving higher values to those who transmitted to the most individuals over shorter and more recent time periods. Specifically, for each individual *u*, we produce a step function representing the number of transmissions from individual *u* over time (Fig. 1) starting from the time of *u*’s first transmission and ending at the termination of the simulation, and we measure the slope of a regression line inferred from the step graph. Because the horizontal axes of all individuals’ step graphs are bounded by the same simulation end time, an individual with more frequent and recent outgoing transmissions with respect to the end-time will have a steeper slope and will thus be assigned a higher value.
- **Metric 3: Indirect Transmissions** This metric expands on Metric 1 in order to quantify an individual’s broader impact on the community. Specifically, for an individual *u*, we count the number of individuals who were infected by somebody who was infected by *u* (i.e., we count the secondary transmissions of *u*).
- **Metric 4: Direct and Indirect Transmissions** This is the sum of Metrics 1 and 3.
- **Metric 5: Number of Contacts** Metric 5 measures each individual’s total number of contacts in the underlying contact network.
- **Metric 6: Number of Contacts and Transmissions** This is the sum of Metrics 1 and 5.

**Figure 1.**
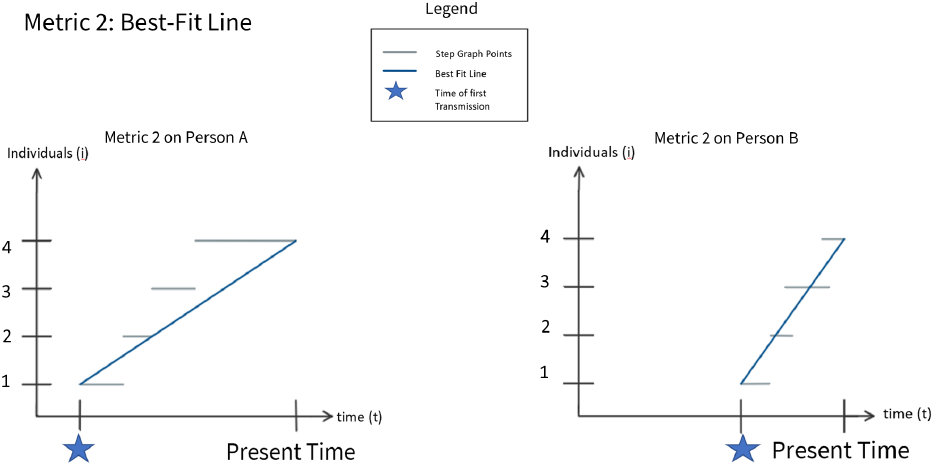
Metric 2. The horizontal axis represents time, and the vertical axis represents the individual’s number of transmissions. Because Person B has transmitted to more individuals more recently than has Person A, the regression line of Person B has a larger slope.

Given a prioritization and the simulated data from which the prioritization was produced, for a given selected metric, SEPIA will compute a value for each individual in the prioritization (Fig. 2). Given a prioritization with a computed metric value for each individual, SEPIA then constructs an “optimal” prioritization by simply sorting the individuals in descending order of metric value. To compare the user-given prioritization against the optimal, SEPIA computes the Kendall Tau-b rank correlation coefficient [7].

**Figure 2.**
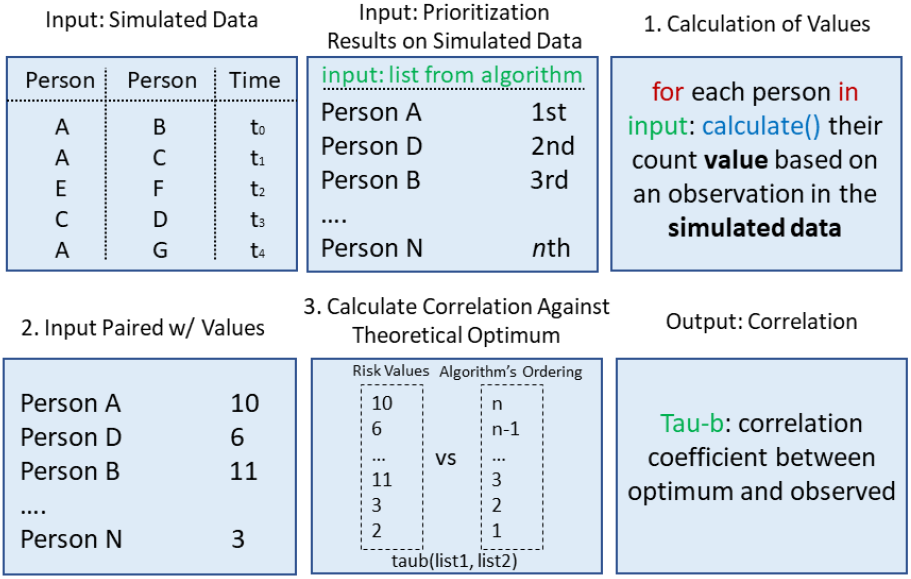
Flowchart of the stages of the SEPIA workflow. The first two boxes indicate SEPIA’s two required inputs: the simulated data and a prioritization of the individuals in the simulated data. In Steps 1 and 2, each individual in the prioritization is paired with a count value, calculated by observing a selected observation (metric) in the simulated data. In Step 3, we calculate the Kendall Tau-b correlation coefficient between these count values and the theoretically optimal ordering of count values.

We used SEPIA to compare the effectiveness of two molecular epidemiological prioritization methods. One approach is to use HIV-TRACE to infer transmission clusters from pairwise distances of viral sequences, monitor the growth of the transmission clusters over time, and prioritize individuals in descending order of transmission cluster growth. The other approach is ProACT [4], a method that utilizes properties of a phylogeny inferred from the viral sequences. We used a simulated dataset produced by FAVITES to emulate the HIV pandemic in San Diego between 2005 and 2014 [5]. The simulated datasets vary the expected degree in the contact network (*E*_*d*_), the rate at which individuals begin ART (λ_+_), and the rate at which individuals stop adhering to ART (λ_−_).

## Results

As can be seen in Figure 3, ProACT consistently out-performed HIV-TRACE transmission cluster growth using all metrics on all simulation conditions. However, both tools consistently had Tau-b scores marginally higher than 0, implying that they are performing only marginally better than a random ordering. As the rate of starting ART (λ_+_) increases, the rate of stopping ART (λ_−_) increases, and the expected degree (*E*_*d*_) increases (i.e., as the outbreak spreads more quickly), ProACT’s performance with respect to metrics 5 and 6 seems to increase slightly. Otherwise, both ProACT and HIV-TRACE transmission cluster growth seem to perform fairly consistently across experimental conditions.

**Figure 3.**
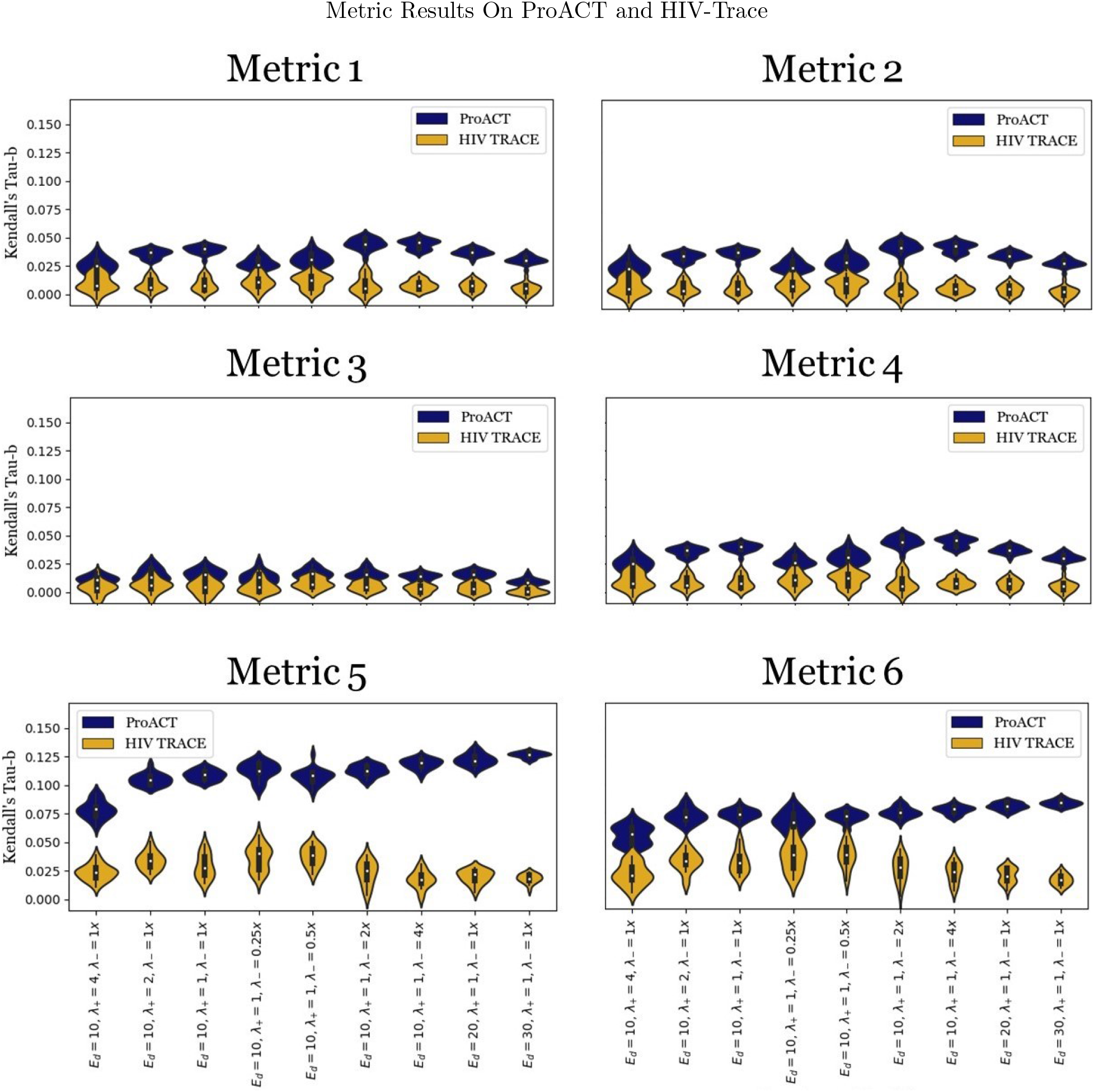
Efficacy of ProACT and HIV-TRACE across all metrics on datasets simulated using FAVITES. The violin plots depict the Kendall Tau-b correlation coefficients between experimental and optimal orderings across 20 replicates for each experimental condition, where *E*_*d*_ denotes the expected number of contacts per individual, λ_+_ denotes the rate at which individuals begin ART, λ_−_ denotes the rate at which individuals stop ART.

## Discussion

Across all defined metrics and all considered simulation conditions, ProACT consistently outperformed prioritization by HIV-TRACE transmission cluster growth. However, both approaches consistently performed just marginally better than a random ordering, implying that there is room for significant improvement in the realm of HIV prioritization.

It must be noted that, while we aimed to provide generalized results by varying key simulation parameters, the simulated epidemics are specifically modeled after the HIV epidemic in San Diego. Molecular epidemiologists will need to assess prioritization techniques using simulated datasets representative of the pathogens and communities in which they are interested.

We hope that SEPIA will enable researchers to quantify and assess the effectiveness of different prioritization approaches in order to select the best existing prioritization method for their communities, develop new prioritization methods that improve upon existing ones, and, ultimately, maximize the impact of the limited resources available to public health officials.

## Author’s contributions

NM conceived and directed this project. All members wrote the code for this project. KA, TJ, ML, TN and MS composed this manuscript.

## Acknowledgements

We would like to acknowledge the Early Research Scholars Program organized by Professor Christine Alvarado and Vignesh Gokul at the University of California, San Diego for granting us this opportunity.

## Availability of data and materials

SEPIA can be found as an open source tool at: https://github.com/Moshiri-Lab/SEPIA

The simulated data can be found at: https://github.com/Moshiri-Lab/SEPIA-paper-final

## Consent for publication

Not Applicable.

## Ethics approval and consent to participate

Not Applicable.

## Author Information

Department of Computer Science and Engineering, University of California San Diego, 9500 Gilman Drive, La Jolla, CA, 92093, USA

Kimberly Almaraz, Tyler Jang, McKenna Lewis, Titan Ngo, Miranda Song and Niema Moshiri

